# Pulling the covers in electronic health records for an association study with self-reported sleep behaviors

**DOI:** 10.1101/341552

**Authors:** Seth D. Rhoades, Lisa Bastarache, Joshua C. Denny, Jacob J. Hughey

## Abstract

The electronic health record (EHR) contains rich histories of clinical care, but has not traditionally been mined for information related to sleep habits. Here we performed a retrospective EHR study and derived a cohort of 3,652 individuals with self-reported sleep behaviors, documented from visits to the sleep clinic. These individuals were obese (mean body mass index 33.6 kg/m^2^) and had a high prevalence of sleep apnea (60.5%), however we found sleep behaviors largely concordant with prior prospective cohort studies. In our cohort, average wake time was one hour later and average sleep duration was 40 minutes longer on weekends than on weekdays (p<1·10^−12^). Sleep duration also varied considerably as a function of age, and tended to be longer in females and in whites. Additionally, through phenome-wide association analyses, we found an association of long weekend sleep with depression, and an unexpectedly large number of associations of long weekday sleep with mental health and neurological disorders (q<0.05). We then sought to replicate previously published genetic associations with morning/evening preference on a subset of our cohort with extant genotyping data (n=555). While those findings did not replicate in our cohort, a polymorphism (rs3754214) in high linkage disequilibrium with a previously published polymorphism near *TARS2* was associated with long sleep duration (p<0.01). Collectively, our results highlight the potential of the EHR for uncovering the correlates of human sleep in real-world populations.

## Introduction

To date, most human sleep research has leveraged prospectively collected cohorts, however many individuals visit their physicians regarding sleep difficulties. Salient information from these visits is captured in the electronic health record (EHR), which opens the possibility of sleep research through retrospective EHR cohorts. EHRs have now matured to a point where longitudinal records exist on large populations, providing an opportunity to drive advances not only in clinical care, but in clinical research (Denny et al. 2016). Moreover, the linking of EHR data with large DNA biobanks is beginning to catalyze scientific discoveries through techniques such as genome-wide and phenome-wide association studies (GWAS and PheWAS) (Denny et al. 2010). Phenome-wide information in the EHR consist of phenotypes produced clinical encounters, such as diagnosis histories or billing codes. This information often receives less attention in clinical sleep research, which typically focuses on pathological sleep conditions in laboratory settings without considering the participant’s medical history (Zee et al. 2014). Additional information outside the clinical setting, such as indications in mental well-being and metabolic health, can otherwise be obtained from self-reports and questionnaires, a common approach in observational sleep research (de Souza & Hidalgo 2014; Konttinen et al. 2014; de Souza & Hidalgo 2015; Vera et al. 2018). These observational studies also typically consider demographic backgrounds, which are additional variables easily obtained from structured medical records. Thus, if we are able to mine sleep-relevant data from the EHR, we can expect to both shed light on new clinical associations with sleep behaviors and corroborate previous sleep studies performed outside an EHR context.

While controlled studies with objective sleep measures provide the strongest evidence for the consequences of sleep disruption, such as the impact on mental health (Minkel et al. 2012), these associations have also been observed through the use of simple questionnaires (Konttinen et al. 2014; Gylen et al. 2014; de Souza & Hidalgo 2015; An et al. 2015). In fact, associations of sleep with age, gender, race, and metabolic parameters such as body mass index (BMI) are largely consistent, regardless of how the sleep metrics are acquired (Ohayon et al. 2004; Lauderdale et al. 2006; Silva et al. 2007; Liu et al. 2012; Rutters et al. 2014; Hashizaki et al. 2015; Fischer et al. 2017; Dietch et al. 2017; Urbanek et al. 2017). Questionnaires such as the Morningness-Eveningness Questionnaire (Horne & Ostberg 1976) and Munich Chronotype Questionnaire (Roenneberg et al. 2003) are common approaches to gauge an individual’s morning or evening preference (i.e., “chronotype”), and are correlated with underlying physiology, including endogenous temperature cycles (Baehr et al. 2000) and dim-light melatonin onset (Kantermann et al. 2015).

Genome-wide association studies on large cohorts (>89,000) from both 23AndMe and the UK Biobank have revealed genetic associations with self-reported chronotype and with sleep duration (Hu et al. 2016; Lane et al. 2016; Jones et al. 2016). Although the overlapping variants across these studies support a causal role for genetics in sleep, the effect sizes tend to be small. In addition, recent work suggests that the influence of genetics on sleep is affected by lifestyle and other behaviors (Vera et al. 2018).

In this work, we explored the potential of the EHR as a resource for clinical sleep research. We first developed a method to extract self-reported sleep behaviors from de-identified EHR at the Vanderbilt University Medical Center. From this method we derived a cohort, and examined associations of their sleep behaviors with demographics, clinical phenotypes, and genetics. Collectively, our results establish the utility of the EHR for retrospective studies of human sleep.

## Materials and Methods

Access to the raw data used in this study is restricted. However, all code and figures related to this study are available at https://figshare.com/s/a614ec7483910e999f9c.

### Sleep phenotype extraction

Our data source is the Synthetic Derivative (SD), Vanderbilt’s database of de-identified medical records (Danciu et al. 2014). To extract sleep behaviors from the SD, we wrote a text parser to detect any mention of sleep in the clinical notes. Within this query, 4,136 notes contained structured fields for “Bedtime on weekdays”, “Bedtime on weekends”, “Wake time on weekdays”, and “Wake time on weekends”. These notes were from visits to the Vanderbilt Sleep Center, and spanned 2002 to 2017 (Fig. S1).

Results from the parser were then manually curated, with 74 notes removed due to vague entries and 177 notes edited due to either parser errors or obvious entry errors (e.g., “12pm” vs. “12am”). Four reports indicating a sleep duration of greater than 18 hours of sleep duration were removed. If an entry contained a time window (e.g., bedtime of 10-11pm), we used the midpoint, unless the range was greater than four hours, in which case we removed the note from the dataset. For this study, we limited the dataset to notes from adults (at least 18 years old at the time of visit to the sleep clinic). Races other than black and white were grouped into “other”. Additionally, we removed non-physiological BMI values greater than 80 and less than 15, which are caused by data entry error. The final dataset consisted of 3,699 reports from 3,652 individuals.

### Statistical analysis

To determine how our sleep cohort compares to a general population from the Vanderbilt EHR, we derived a 1:1 matched cohort in the SD by age (in years, at the time of their last visit in the records), sex, race, and duration of record (in years). Pairwise comparisons across cohorts were performed using student’s t-tests, and prevalence of phenotype codes through a two-proportions z-test. To account for the cyclical nature of clock time, we adjusted bed, wake, and midpoint times as a difference in hours from the circular mean. We calculated additional quantitative phenotypes from the sleep self-reports, including sleep duration and measures of weekday-to-weekend shifts in sleep behaviors (called “social jetlag” or “social sleep lag” in the sleep literature (Wittmann et al. 2006)). Social sleep lag was calculated by the circular difference between weekday and weekend sleep midpoints, positive if individuals delayed their weekend midpoint, and negative if individuals advanced their weekend midpoint. Pairwise comparisons of sleep behaviors within the sleep cohort, such as sleep duration by gender, were also performed using student’s t-tests.

We modeled sleep behaviors as a function of demographic variables and BMI at the time of the sleep clinic visit using an ordinal regression approach with the rms R package v5.1-2 (Harrell 2018). BMI values were log-transformed before modeling, and age was fit as a restricted spline. Model selection consisted of comparing coefficients across a variable number of knots in the restricted splines, and comparing model fits using likelihood ratio tests. Both of these methods jointly determined the flexibility of the fits to age and the justification of higher-order terms in the model. We performed analysis in R v3.4.1, and generated plots using ggplot2 v2.2.1.

### Phenome-wide association analysis

We explored associations between self-reported sleep behaviors and phenotype codes (“phecodes”), which have been mapped to related ICD-9-CM codes for research purposes. Details for these mappings are described elsewhere (Denny et al. 2013). These clinical diagnoses were modeled as dependent variables, with sleep duration or social sleep lag as independent variables in a generalized linear model, and age, gender, and race were included as covariates. Cases for a particular phecode consisted of subjects with that phecode in the record on at least two distinct dates, whereas controls had zero instances of the respective phecode. Each phecode defines a control group for analysis by using a set of exclusion phecodes (based on version 1.2 of the phecode mappings available at http://phewascatalog.org (Denny et al. 2013)). Thus individuals who do not have the phecode of interest but have an exclusion phecode are considered neither cases nor controls and removed from the model. We analyzed only those phecodes with a prevalence of at least 1% in the sleep cohort, and accounted for multiple-testing through a false-discovery rate procedure (Benjamini & Hochberg 1995).

### Genetic association analysis

To find genetic associations with sleep phenotypes, we leveraged multiple data sources within BioVU, Vanderbilt’s de-identified DNA biobank linked to the SD (Roden et al. 2008). Genotyping data comes from the Illumina Infinium Human Exome BeadChip (Cronin et al. 2014) and the Illumina Infinium Expanded Multi-Ethnic Genotyping Array (MEGA^EX^). We considered data on individuals of European ancestry. 111 individuals in the sleep cohort had data on both platforms, and for any variant discrepancies across the two platforms, we used the calls from the Exome BeadChip. Collectively, we performed genetic association analysis on 555 unique individuals in the sleep cohort. We compiled a list of SNPs having significant associations with self-reported chronotype (Hu et al. 2016; Lane et al. 2016; Jones et al. 2016), and expanded our search by considering tagging SNPs that are in high LD (r^2^ > 0.80) with the published SNPs. We identified tagging SNPs on European ancestry genotype data from Phase 3 (version 5) of the 1000 Genomes Project using the LDproxy tool at https://analysistools.nci.nih.gov/LDlink/. SNPs with less than 1% minor allelic frequency (MAF) were removed from consideration. Associations with sleep phenotypes were modeled by ordinal regression, with additive genetic effects and adjustments for age and gender. We performed power analysis using the method of (Derkach et al. 2018), with inputs of MAF and effect size from the GWAS results on a continuous chronotype measurement (Lane et al. 2016; Jones et al. 2016).

## Results

### Characteristics of a sleep cohort obtained from the EHR

We searched the clinical notes in the Vanderbilt Synthetic Derivative (SD) for mentions of sleep behavior, and found that notes from the sleep clinic often contain structured information on patients’ self-reported bedtimes and wake-times on weekdays and weekends (see Materials and Methods for details, Fig. S1). We parsed these notes to yield a dataset of 3,699 sleep reports from 3,652 individuals (which we call the sleep cohort, Fig. S2). This cohort consists of predominantly white adults (Table 1), which is reflective of the typical population in the Vanderbilt SD, and is also generally obese, with subjects having a BMI of 33.6 ± 8.9 kg /m^2^ (mean ± S.D.) at the time of their visit to the sleep clinic.

**Table 1.** Cohort characteristics. The sleep cohort was matched by length of record, age, gender, and race. Numbers for gender, race, and ethnicity correspond to percentages. Numbers for age, phecodes, and visits correspond to mean (SD). P-values are based on a student’s t-test.

|  | Sleep Cohort (n=3652) | Matched Cohort (n=3646) | P-value |
| --- | --- | --- | --- |
| Gender (male / female) | 50.7 / 49.3 | - | - |
| Race (black / white / other) | 14.2 / 80.6 / 5.2 | - | - |
| Age at visit | 48.0 (13.7) | - | - |
| Total number of unique phecodes | 54.0 (52.1) | 27.7 (30.6) | $2.9 \cdot 10^{-146}$ |
| Number of visits | 91.3 (111.1) | 38.8 (60.4) | $5.3 \cdot 10^{-133}$ |

To assess the clinical features of this cohort, we calculated the prevalence of diagnoses in their records (based on phecodes, a grouping of ICD-9 codes designed for high-throughput analysis (Denny et al. 2010)), and matched a cohort in the SD based on age, sex, race, and duration of the medical record. The diagnosis with the highest prevalence in the sleep cohort corresponds to obstructive sleep apnea (60.5%, compared to 5.3% in the matched cohort, Fig. S3). Other highly prevalent phecodes in the sleep cohort included known comorbidities of obstructive sleep apnea (Somers et al. 2008), such as obesity, hypertension, and hyperlipidemia (p<1·10^−56^ by two-proportions z-test). Additionally, the number of total clinical encounters differed significantly between cohorts (Table 1), indicating that the individuals visiting the sleep clinic are heavy users of the healthcare system, and not necessarily representative of healthy adults. Nonetheless, the sleep cohort presents an opportunity to examine the correlates of self-reported sleep behaviors in a real-world population.

### Relationships between EHR-derived, self-reported sleep behaviors and demographics in the sleep cohort

We next examined the distributions of self-reported bedtimes and wake-times on weekdays and weekends, and their relationships with gender, race, and age. As expected, we observed large shifts between weekdays and weekends. Mean weekday and weekend wake-times shifted from 6:20 a.m. to 7:20 a.m., respectively (Fig. 1, p=1.86·10^−88^), and sleep midpoint shifted from 2:22 a.m. on weekdays to 3:03 a.m. on weekends (p=4.52·10^−13^).

**Figure 1.**
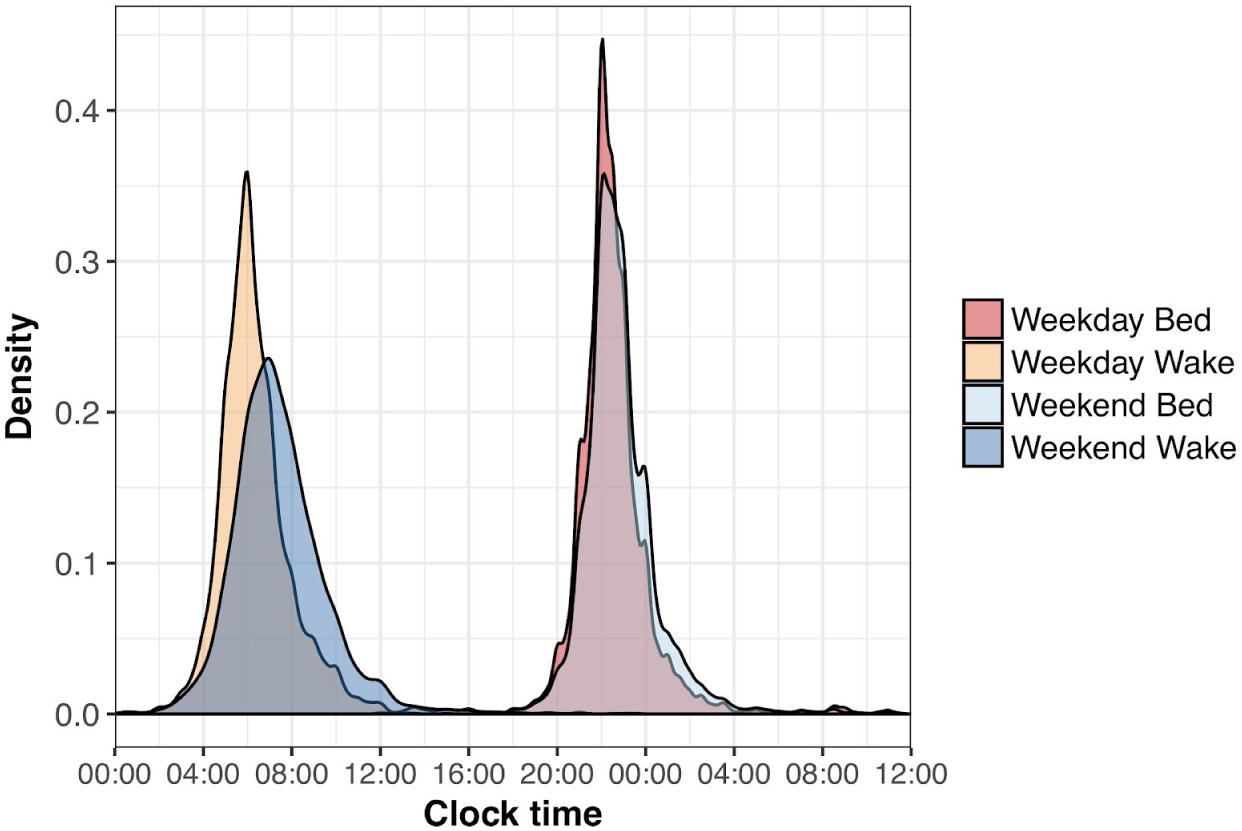
Distribution of self-reported bed and wake times for the sleep cohort. Bed times past midnight were adjusted to separate bed and wake times and ease visualization.

Both weekday and weekend sleep durations were associated with gender, race, and age in an ordinal regression model (p<0.05, Table 2, Fig. 2, Supplementary Files 1-2). Specifically, sleep durations tended to be longer in females and whites, which has been observed in prior studies (Lauderdale et al. 2006; Dietch et al. 2017). Weekday-to-weekend sleep midpoint shift, i.e. “social sleep lag”, also expectedly varied across age (Table 2, Fig. 3A, Supplementary File 3), demonstrating the extent to which younger individuals shift their sleep schedules (Hashizaki et al. 2015; Koopman et al. 2017). Social sleep lag, however, was not significantly associated with gender or race. Weekend sleep midpoint also associated strongly with age, although race and gender showed little effect (Table 2, Supplementary File 4). As metabolic health has been closely tied to sleep behaviors (Liu et al. 2012; Reutrakul et al. 2013; Rutters et al. 2014), we also checked for associations of sleep midpoint with BMI at the time of visit to the sleep clinic. BMI, and interactions between BMI and gender, displayed strong associations with sleep midpoint, in addition to interactions between gender and age (Table 2, Fig. 3B). Conditional on the model, heavier individuals were predicted to continually shift their sleep midpoint later with age, whereas thinner individuals were predicted to maintain relatively stable midpoints in adulthood. These associations did not hold for weekday sleep midpoint, with age as the only significant covariate (not shown). We also did not find a significant association of BMI in any other sleep phenotype model (Table 2). Collectively, the relationships between these EHR-derived, self-reported sleep behaviors and demographic variables demonstrate a general concordance with previous studies.

**Figure 2.**
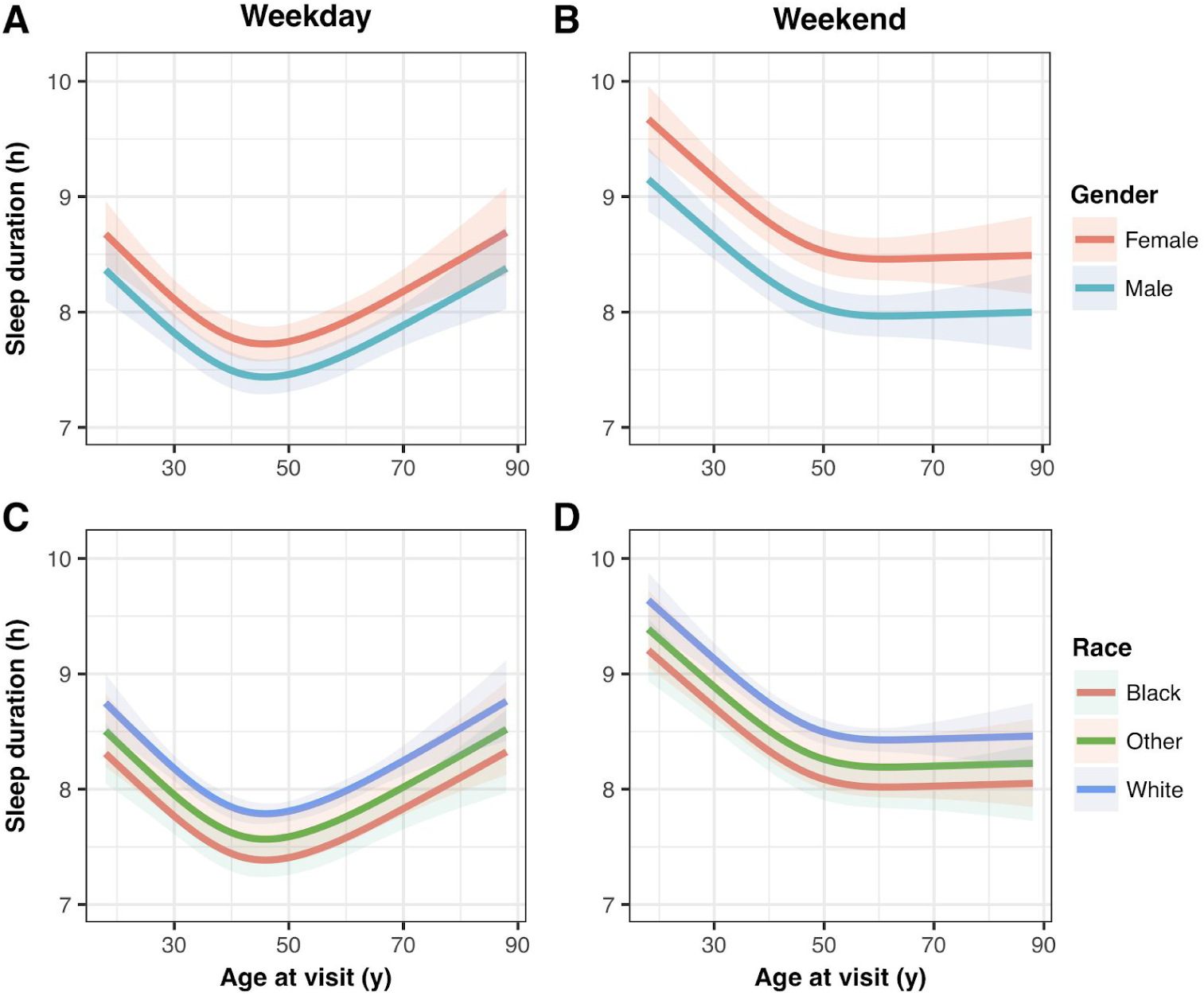
Weekday and weekend sleep duration (in hours) as a function of age across gender (A, B) and race (C, D) in the sleep cohort. Gender, and race, and age as a restricted spline were all significant predictor variables in both weekday and weekend models. Shaded regions represent best-fit 95% confidence intervals in an ordinal regression procedure.

**Figure 3.**
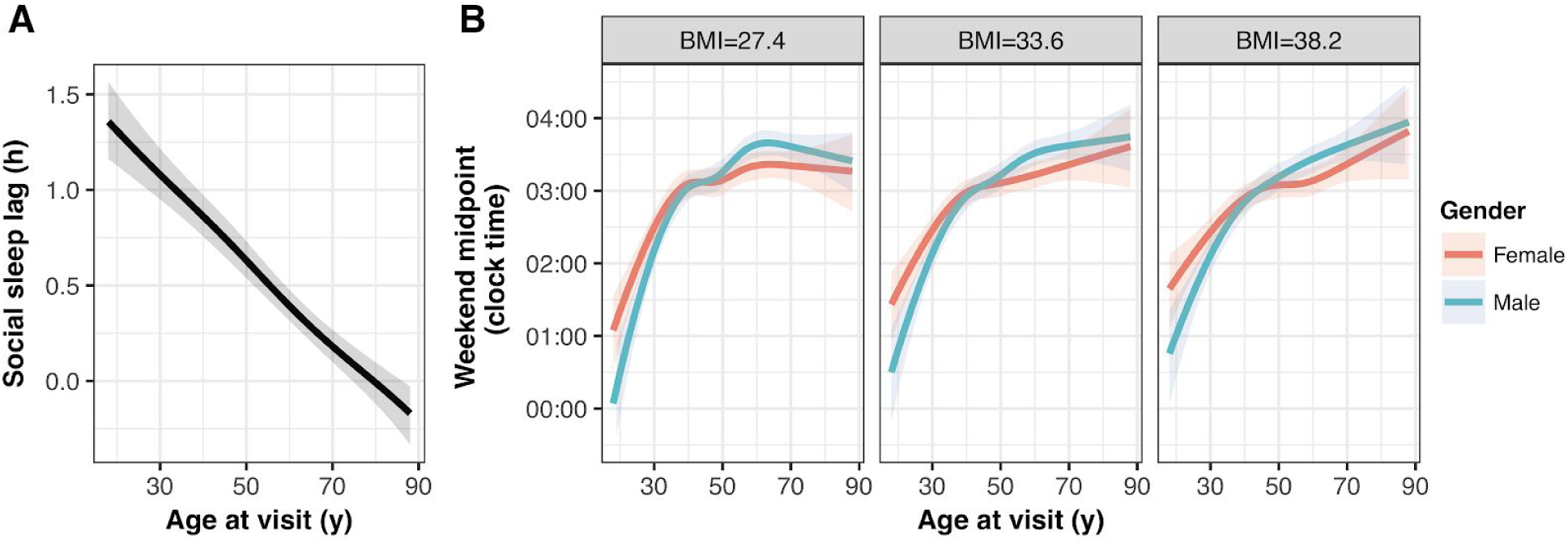
(A) Social sleep lag (in hours) as a function of age as a restricted spline in the sleep cohort. Gender nor race were significant predictors in this model, and thus the model predictions for the entire cohort is depicted here. (B) Weekend sleep midpoint, adjusted to the circular mean of midpoint for the sleep cohort, with model predictions across gender and the 1st quartile, mean, and 3rd quartile of BMI, which were significant interaction variables with age in the chosen model. Shaded regions represent best-fit 95% confidence intervals in an ordinal regression procedure.

**Table 2.** Model statistics for final weekday sleep duration (n=3405), weekend sleep duration (n=3399), social sleep lag (n=3324), and weekend sleep midpoint (n=3391) models in an ordinal regression procedure. Numbers represent p-values for each predictor variable, and asterisks indicate the model described in the main text and figures. Age is modeled as a restricted spline in both individual and interaction terms which contain age. Model building and selection is detailed in Supplementary Files 1-4.

|  | Gender | Race | Age | BMI | Age * BMI | Age * Gender |
| --- | --- | --- | --- | --- | --- | --- |
| **Weekday sleep duration** |  |  |  |  |  |  |
| Model 1 | $8.65 \cdot 10^{-8}$ | - | - | - | - | - |
| Model 2 | $2.97 \cdot 10^{-9}$ | $8.24 \cdot 10^{-8}$ | - | - | - | - |
| Model 3* | $1.81 \cdot 10^{-9}$ | $5.81 \cdot 10^{-8}$ | $3.16 \cdot 10^{-17}$ | - | - | - |
| Model 4 | $6.88 \cdot 10^{-10}$ | $4.47 \cdot 10^{-7}$ | $1.36 \cdot 10^{-15}$ | 0.10 | - | - |
| **Weekend sleep duration** |  |  |  |  |  |  |
| Model 1 | $4.88 \cdot 10^{-17}$ | - | - | - | - | - |
| Model 2 | $2.24 \cdot 10^{-18}$ | $3.34 \cdot 10^{-4}$ | - | - | - | - |
| Model 3* | $1.55 \cdot 10^{-17}$ | $6.49 \cdot 10^{-6}$ | $4.72 \cdot 10^{-22}$ | - | - | - |
| Model 4 | $1.58 \cdot 10^{-17}$ | $8.06 \cdot 10^{-6}$ | $5.92 \cdot 10^{-22}$ | 0.80 | - | - |
| **Social sleep lag** |  |  |  |  |  |  |
| Model 1 | $3.46 \cdot 10^{-3}$ | - | - | - | - | - |
| Model 2 | $5.38 \cdot 10^{-3}$ | 0.03 | - | - | - | - |
| Model 3* | 0.08 | 0.87 | $6.59 \cdot 10^{-76}$ | - | - | - |
| Model 4 | 0.09 | 0.86 | $1.26 \cdot 10^{-75}$ | 0.87 | - | - |
| **Weekend sleep midpoint** |  |  |  |  |  |  |
| Model 1 | 0.01 | - | - | - | - | - |
| Model 2 | 0.02 | 0.29 | - | - | - | - |
| Model 3 | 0.11 | 0.77 | $4.02 \cdot 10^{-80}$ | - | - | - |
| Model 4 | 0.16 | 0.56 | $2.09 \cdot 10^{-80}$ | $8.56 \cdot 10^{-3}$ | - | - |
| Model 5 | 0.23 | 0.60 | $1.55 \cdot 10^{-80}$ | $8.44 \cdot 10^{-3}$ | $5.42 \cdot 10^{-4}$ | - |
| Model 6* | 0.24 | 0.58 | $9.32 \cdot 10^{-81}$ | $7.96 \cdot 10^{-3}$ | $2.21 \cdot 10^{-4}$ | $5.18 \cdot 10^{-5}$ |

### Associations between sleep behaviors and clinical phenotypes

We then analyzed the extent to which our cohort’s sleep behaviors were associated with clinical diagnoses (Denny et al. 2010), adjusting for gender, age, and race. In this PheWAS approach, we found that longer self-reported sleep duration on weekends most strongly associated with depression (q = 2.49·10^−3^, Fig. 4A), which is consistent with a recent meta-analysis (Zhai et al. 2015). Sleep midpoint on weekends, on the other hand, was not significantly associated with any clinical phenotypes. Unexpectedly, sleep duration on weekdays was associated with a large number of phenotypes, including many mental and neurological disorders (Fig. 4B). These phenotypes may covary in the sleep cohort, which could explain the lower-than-expected test statistics of the quantile-quantile plot (Fig. S4). Overall, we found few instances in which a higher prevalence of the clinical phenotype was significantly associated with shorter sleep, which may be a result of our cohort containing few short sleepers. Individuals who advance their sleep schedules on weekends demonstrated an increased association with respiratory and neurological diagnoses (Fig. 4C). Taken together, these results replicate previous associations between sleep and mental health and suggest new hypotheses for future investigation.

**Figure 4.**
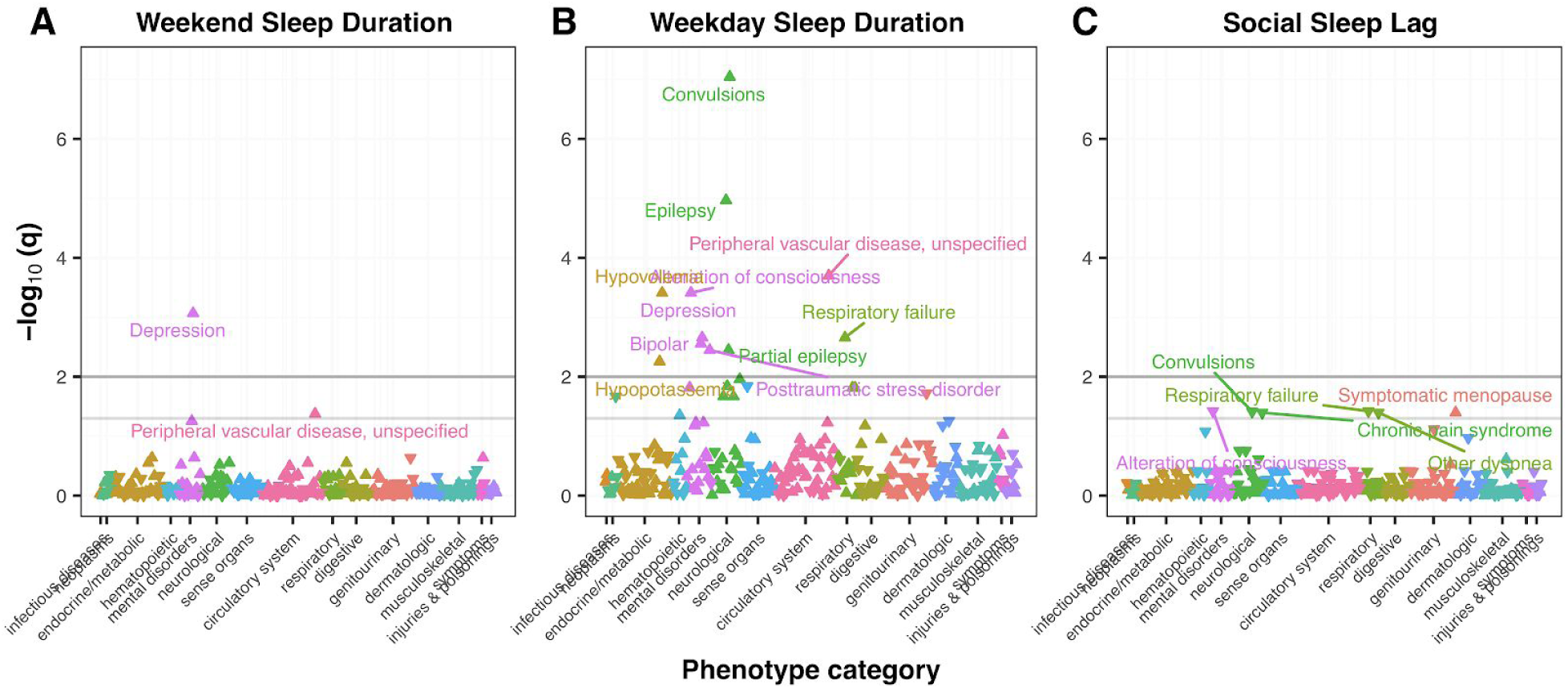
PheWAS for (A) weekend sleep duration and (B) weekday sleep duration. Light grey lines indicate q=0.05, and darker grey lines q=0.01. Phecodes with q<0.05 are annotated, except for weekday sleep duration (for ease of visualization).

### Targeted replication of associations between sleep behaviors and genetic variants

Of the 3,647 individuals in the sleep cohort, 555 have genotype data available through BioVU, Vanderbilt’s de-identified DNA biobank linked to the SD (Roden et al. 2008). Because of this relatively small size, rather than performing a genome-wide search for associations between sleep behaviors and genetic variation, we instead attempted to replicate significant associations between single-nucleotide polymorphisms (SNPs) and self-reported morningness-eveningness from much larger recent sleep GWASes (Hu et al. 2016; Lane et al. 2016; Jones et al. 2016). Three such SNPs from these studies are assayed on our genotyping platform and have a mean allelic frequency greater than 1%: rs12140153, rs1144566, and rs35333999. However, our power to detect associations with these SNPs was 0.51, 0.43, and 0.38, respectively, which likely explains a lack of association with all of our sleep phenotypes. We expanded our search by considering two additional SNPs, rs3754214 and rs9753974, in high linkage disequilibrium with the published SNPs. Of these, rs3754214, close to rs10157197 (located near *TARS2*, r^2^= 0.90), was associated with increased weekday (β = 0.44 (0.12), p=1.54·10^−4^, Fig. 5A) and weekend sleep duration (β = 0.31 (0.11), p=0.006, Fig. 5B, Table 3). Our data shows greater than 15 minutes in increased sleep duration for each C allele, which far exceeds effect sizes of SNPs found to associate with sleep duration (Jones et al. 2016). While rs10157197 is one of thirteen SNPs whose associations replicated across the 23AndMe and UK Biobank datasets for chronotype (Hu et al. 2016; Jones et al. 2016), Jones et al. did not find such an association with sleep duration. Although these results are preliminary, they suggest that the previously observed genetic contributions to sleep may be moderated by the health-related characteristics of individuals in the sleep cohort.

**Figure 5.**
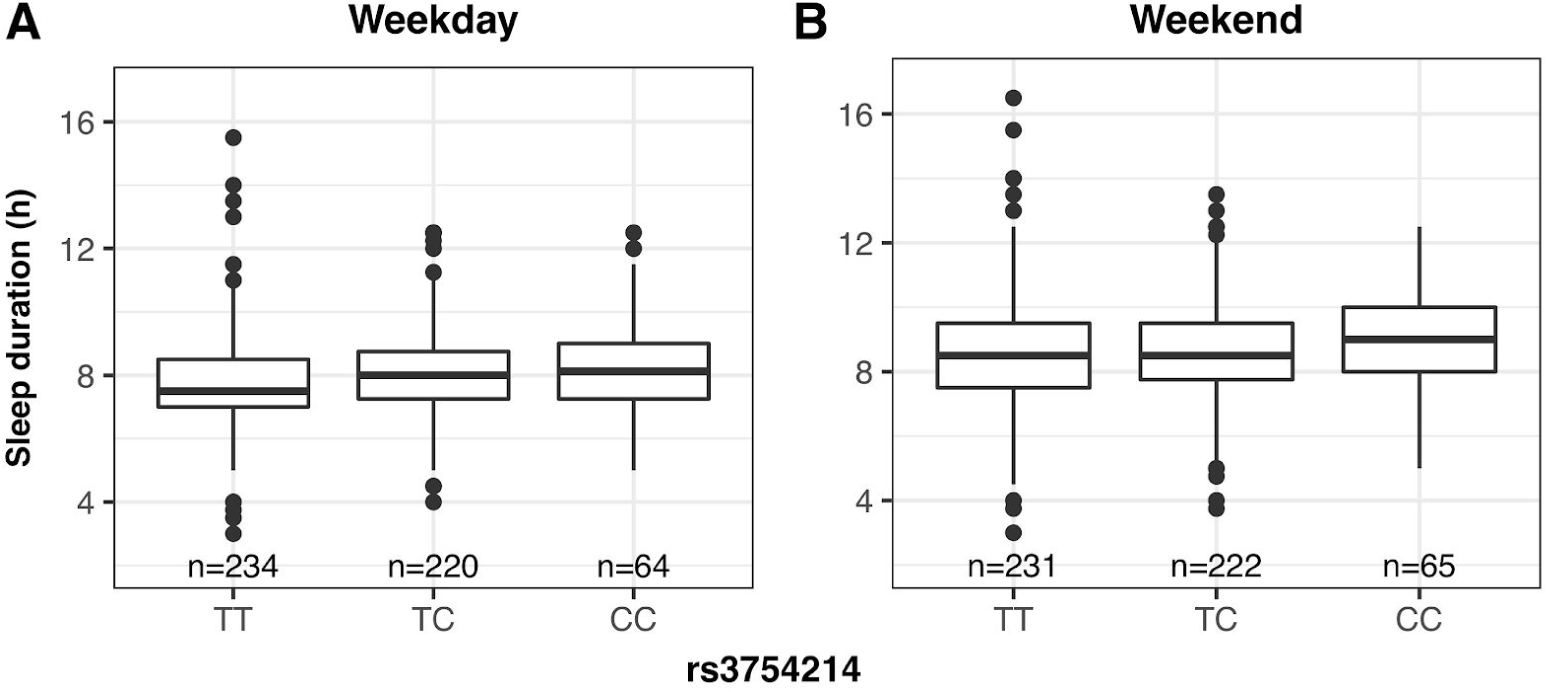
(A) Weekend sleep duration and (B) weekday sleep duration by alleles of variant rs3754214 in the sleep cohort with extant genotyping (n=555), significant in an ordinal regression procedure.

**Table 3.** Genetic association analysis with weekday and weekend sleep duration in an ordinal regression model (n=555). Asterisks indicate SNPs previously associated with chronotype, while those without astericks indicate SNPs in high LD (r^2^ > 0.80) with any of these published SNPs.

|  | Coef | S.E. | P-value |
| --- | --- | --- | --- |
| **Weekday sleep duration** |  |  |  |
| rs3754214 | 0.44 | 0.12 | $1.54 \cdot 10^{-4}$ |
| rs9573974 | 0.48 | 0.29 | 0.11 |
| rs35333999* | 0.36 | 0.35 | 0.31 |
| rs1144566* | 0.14 | 0.30 | 0.64 |
| rs12140153* | 0.10 | 0.21 | 0.66 |
| **Weekend sleep duration** |  |  |  |
| rs3754214 | 0.31 | 0.11 | $6.21 \cdot 10^{-3}$ |
| rs9573974 | 0.59 | 0.29 | 0.04 |
| rs35333999* | -0.49 | 0.30 | 0.11 |
| rs1144566* | -0.05 | 0.35 | 0.88 |
| rs12140153* | -0.02 | 0.21 | 0.91 |

## Discussion

In this study, we parsed notes in the EHR for any mention of sleep and discovered structured entries for self-reported bed and wake times at the sleep clinic. Although the cohort is not representative of healthy adults, we found associations of the sleep patterns comparable to recent work, establishing the suitability of this dataset for exploratory analysis of sleep behaviors with phenome-wide and genetic information.

The sleep behaviors used in this study come from routine questions asked by the physician at the sleep clinic, which raises several limitations. These questions are not derived from validated sleep questionnaires, and many of the responses regarding bed and wake times were imprecise. Beyond concerns of precision, we cannot be sure of the extent of bias in how patients respond to clinicians’ questions compared to how they would respond to a validated questionnaire. Finally, the generalizability of our approach depends on the extent to which similar information is obtainable in other institutions’ EHRs. Future work may benefit from more sophisticated natural language processing techniques to identify mentions of sleep-related behaviors (such as shift work) in unstructured text outside sleep clinic notes.

The observational nature of the EHR makes determination of causality difficult. Furthermore, most individuals in our cohort have only one clinical encounter with sleep information, and the phenotypes in our phenome-wide association study are based on each subject’s entire record, both before and after the sleep clinic visit(s). We expect that integration of the EHR with longitudinal sleep assessments, such as from wearables and continuous positive airway pressure ventilators, will help us unravel the time-dependent relationships between sleep and other clinical phenotypes (Hwang 2016; Baron et al. 2017).

Although this study is based on a convenience sample, many of our findings are consistent with previous studies based on prospective cohorts. For example, weekday sleep duration follows a U-shaped curve as a function of age, and ultimately converges with weekend sleep duration in older individuals. These patterns are consistent with expected constraints of working-age adults and altered sleep requirements in the elderly, and have been observed previously (Ohayon et al. 2004; Silva et al. 2007; Hashizaki et al. 2015). Both sleep midpoint and sleep lag also vary as a function of age, which has been observed in adolescents and young adults (Rutters et al. 2014; Hashizaki et al. 2015; Fischer et al. 2017; Koopman et al. 2017; Urbanek et al. 2017). Gender and race did not explain social sleep lag, which may reflect broader cultural norms and chronotype shifts in younger individuals regardless of background. The average weekend sleep duration of 8.59 hours closely matches sleep durations from other studies based on self-reported data (Liu et al. 2012; Rutters et al. 2014; Fischer et al. 2017; Koopman et al. 2017), which commonly overestimates sleep duration compared to objective measures such as actigraphy (Lauderdale et al. 2006; Silva et al. 2007; Arora et al. 2013; Dietch et al. 2017). Well-established trends in both objective and subjective sleep analyses demonstrate whites sleep upwards of 45 minutes longer than blacks (Lauderdale et al. 2006; Dietch et al. 2017), and women sleep upwards of 30 minutes longer than men (Lauderdale et al. 2006; Liu et al. 2012; Dietch et al. 2017; Urbanek et al. 2017), which are all consistent with our findings.

We find a greater discordance to prior work with our cohort’s sleep midpoints, which typically vary from from 3:00 a.m. to 4:30 a.m. in adults (Lucassen et al. 2013; Rutters et al. 2014; Hashizaki et al. 2015; Fischer et al. 2017). Our cohort’s sleep midpoints resemble those of morning chronotypes among obese adults (Lucassen et al. 2013) and elderly individuals (Fischer et al. 2017), which may speak to a morning preference in our older and obese cohort. Although multiple studies have investigated interactions between metabolic health and sleep (Liu et al. 2012; Reutrakul et al. 2013; Rutters et al. 2014), our cohort shows little evidence of an association between BMI and sleep duration, which may be a consequence of our cohort’s high prevalence of obesity. Our analysis did identify a moderate effect of BMI on sleep midpoint in older individuals, highlighting a potential metabolic influence on sleep during aging.

Deriving sleep behaviors from the EHR allowed us to quantify associations with a broad range of clinically-defined phenotypes. The association between long weekend sleep and depression is supported by questionnaire-based studies (Sun et al. 2018), however, long weekday sleep duration unexpectedly shows very strong associations with clusters of mental health and neurological conditions. One possible explanation for the differences between PheWAS analyses for sleep duration on weekdays and weekends is that long sleep duration on weekends stems from accumulating sleep debt during the week, while abnormal sleep patterns on weekdays may occur despite social and societal constraints and have a deeper clinical basis. Given our cohort consists of relatively fewer short sleepers, associations of disease with short sleep noted in the literature are likely muted. Nonetheless our findings could open new avenues of research to fully understand the risks of excessive sleep.

The limited number of individuals in the sleep cohort with genotype data available prevented a thorough attempt to replicate known genetic associations. Only one prior GWAS to date analyzed both chronotypes and sleep duration (Jones et al. 2016). The variants found to associate with sleep duration were not included in our exome array, effectively limiting our genetic association analysis to variants associated with chronotype. Thus our power analysis is only approximate, as the published effect sizes for chronotype may not align with our sleep phenotypes. Even so, we are still likely underpowered to detect these associations in our cohort. Although we identified a variant in high LD with rs10157197 that was associated with sleep duration, the effect size is considerably larger than anything observed in the previous GWAS. Additional studies with larger sample sizes will be needed to further delineate these effects.

In conclusion, our findings support the use of the EHR for sleep research on clinically relevant populations. As the EHR grows to include data from consumer devices that monitor sleep and other behaviors, approaches like ours may help reveal the relationships between sleep and other aspects of human health on an unprecedented scale, and we expect advances in clinical informatics will continue to benefit sleep research.

## Acknowledgments

We thank the Vanderbilt Biostatistics Clinic for suggestions on the statistical modeling, and Jamie Andry for insight on clinical notes from the Vanderbilt Sleep Center. The MEGA^EX^ genotyping data was obtained from Vanderbilt University Medical Center’s BioVU which is supported by institutional funding, the 1S10RR025141-01 instrumentation award, and by the CTSA grant UL1TR000445 from NCATS/NIH, with additional funding provided by the NIH through grants P50GM115305 and U19HL065962. SDR is supported by NIH T32HG008341, JCD by NIH R01LM016085, JJH by NIH R35GM124685.

---

**Supplementary File 1.** Model selection process, and final model summaries, for weekday sleep duration.

**Supplementary File 2.** Model selection process, and final model summaries, for weekend sleep duration.

**Supplementary File 3.** Model selection process, and final model summaries, for social sleep lag.

**Supplementary File 4.** Model selection process, and final model summaries, for weekend sleep midpoint.

## References

An KO, Jang JY, Kim J. 2015. Sedentary Behavior and Sleep Duration Are Associated with Both Stress Symptoms and Suicidal Thoughts in Korean Adults. Tohoku J Exp Med. 237:279–286.

Arora T, Broglia E, Pushpakumar D, Lodhi T, Taheri S. 2013. An investigation into the strength of the association and agreement levels between subjective and objective sleep duration in adolescents. PLoS One. 8:e72406.

Baehr EK, Revelle W, Eastman CI. 2000. Individual differences in the phase and amplitude of the human circadian temperature rhythm: with an emphasis on morningness-eveningness. J Sleep Res. 9:117–127.

Baron KG, Duffecy J, Berendsen MA, Cheung Mason I, Lattie EG, Manalo NC. 2017. Feeling validated yet? A scoping review of the use of consumer-targeted wearable and mobile technology to measure and improve sleep. Sleep Med Rev [Internet]. Available from: http://dx.doi.org/10.1016/j.smrv.2017.12.002

Benjamini Y, Hochberg Y. 1995. Controlling the False Discovery Rate: A Practical and Powerful Approach to Multiple Testing. J R Stat Soc Series B Stat Methodol. 57:289–300.

Cronin RM, Field JR, Bradford Y, Shaffer CM, Carroll RJ, Mosley JD, Bastarache L, Edwards TL, Hebbring SJ, Lin S, et al. 2014. Phenome-wide association studies demonstrating pleiotropy of genetic variants within FTO with and without adjustment for body mass index. Front Genet. 5:250.

Danciu I, Cowan JD, Basford M, Wang X, Saip A, Osgood S, Shirey-Rice J, Kirby J, Harris PA. 2014. Secondary use of clinical data: the Vanderbilt approach. J Biomed Inform. 52:28–35.

Denny JC, Bastarache L, Ritchie MD, Carroll RJ, Zink R, Mosley JD, Field JR, Pulley JM, Ramirez AH, Bowton E, et al. 2013. Systematic comparison of phenome-wide association study of electronic medical record data and genome-wide association study data. Nat Biotechnol. 31:1102–1110.

Denny JC, Bastarache L, Roden DM. 2016. Phenome-Wide Association Studies as a Tool to Advance Precision Medicine. Annu Rev Genomics Hum Genet. 17:353–373.

Denny JC, Ritchie MD, Basford MA, Pulley JM, Bastarache L, Brown-Gentry K, Wang D, Masys DR, Roden DM, Crawford DC. 2010. PheWAS: demonstrating the feasibility of a phenome-wide scan to discover gene-disease associations. Bioinformatics. 26:1205–1210.

Derkach A, Zhang H, Chatterjee N. 2018. Power Analysis for Genetic Association Test (PAGEANT) provides insights to challenges for rare variant association studies. Bioinformatics. 34:1506–1513.

Dietch JR, Taylor DJ, Smyth JM, Ahn C, Smith TW, Uchino BN, Allison M, Ruiz JM. 2017. Gender and racial/ethnic differences in sleep duration in the North Texas heart study. Sleep Health. 3:324–327.

Fischer D, Lombardi DA, Marucci-Wellman H, Roenneberg T. 2017. Chronotypes in the US - Influence of age and sex. PLoS One. 12:e0178782.

Gylen E, Anttalainen U, Saaresranta T. 2014. Relationship between habitual sleep duration, obesity and depressive symptoms in patients with sleep apnoea. Obes Res Clin Pract. 8:e459–65.

Harrell FE Jr. 2018. rms: Regression Modeling Strategies. R package version 5.1-2 [Internet]. [place unknown]. Available from: https://CRAN.R-project.org/package=rms

Hashizaki M, Nakajima H, Kume K. 2015. Monitoring of Weekly Sleep Pattern Variations at Home with a Contactless Biomotion Sensor. Sensors. 15:18950–18964.

Horne JA, Ostberg O. 1976. A self-assessment questionnaire to determine morningness-eveningness in human circadian rhythms. Int J Chronobiol. 4:97–110.

Hu Y, Shmygelska A, Tran D, Eriksson N, Tung JY, Hinds DA. 2016. GWAS of 89,283 individuals identifies genetic variants associated with self-reporting of being a morning person. Nat Commun. 7:10448.

Hwang D. 2016. Monitoring Progress and Adherence with Positive Airway Pressure Therapy for Obstructive Sleep Apnea: The Roles of Telemedicine and Mobile Health Applications. Sleep Med Clin. 11:161–171.

Jones SE, Tyrrell J, Wood AR, Beaumont RN, Ruth KS, Tuke MA, Yaghootkar H, Hu Y, Teder-Laving M, Hayward C, et al. 2016. Genome-Wide Association Analyses in 128,266 Individuals Identifies New Morningness and Sleep Duration Loci. PLoS Genet. 12:e1006125.

Kantermann T, Sung H, Burgess HJ. 2015. Comparing the Morningness-Eveningness Questionnaire and Munich ChronoType Questionnaire to the Dim Light Melatonin Onset. J Biol Rhythms. 30:449–453.

Konttinen H, Kronholm E, Partonen T, Kanerva N, Männistö S, Haukkala A. 2014. Morningness-eveningness, depressive symptoms, and emotional eating: a population-based study. Chronobiol Int. 31:554–563.

Koopman ADM, Rauh SP, van ’t Riet E, Groeneveld L, van der Heijden AA, Elders PJ, Dekker JM, Nijpels G, Beulens JW, Rutters F. 2017. The Association between Social Jetlag, the Metabolic Syndrome, and Type 2 Diabetes Mellitus in the General Population: The New Hoorn Study. J Biol Rhythms. 32:359–368.

Lane JM, Vlasac I, Anderson SG, Kyle SD, Dixon WG, Bechtold DA, Gill S, Little MA, Luik A, Loudon A, et al. 2016. Genome-wide association analysis identifies novel loci for chronotype in 100,420 individuals from the UK Biobank. Nat Commun. 7:10889.

Lauderdale DS, Knutson KL, Yan LL, Rathouz PJ, Hulley SB, Sidney S, Liu K. 2006. Objectively measured sleep characteristics among early-middle-aged adults: the CARDIA study. Am J Epidemiol. 164:5–16.

Liu R, Liu X, Arguelles LM, Patwari PP, Zee PC, Chervin RD, Ouyang F, Christoffel KK, Zhang S, Hong X, et al. 2012. A population-based twin study on sleep duration and body composition. Obesity. 20:192–199.

Lucassen EA, Zhao X, Rother KI, Mattingly MS, Courville AB, de Jonge L, Csako G, Cizza G, Sleep Extension Study Group. 2013. Evening chronotype is associated with changes in eating behavior, more sleep apnea, and increased stress hormones in short sleeping obese individuals. PLoS One. 8:e56519.

Minkel JD, Banks S, Htaik O, Moreta MC, Jones CW, McGlinchey EL, Simpson NS, Dinges DF. 2012. Sleep deprivation and stressors: evidence for elevated negative affect in response to mild stressors when sleep deprived. Emotion. 12:1015–1020.

Ohayon MM, Carskadon MA, Guilleminault C, Vitiello MV. 2004. Meta-analysis of quantitative sleep parameters from childhood to old age in healthy individuals: developing normative sleep values across the human lifespan. Sleep. 27:1255–1273.

Reutrakul S, Hood MM, Crowley SJ, Morgan MK, Teodori M, Knutson KL, Van Cauter E. 2013. Chronotype is independently associated with glycemic control in type 2 diabetes. Diabetes Care. 36:2523–2529.

Roden DM, Pulley JM, Basford MA, Bernard GR, Clayton EW, Balser JR, Masys DR. 2008. Development of a large-scale de-identified DNA biobank to enable personalized medicine. Clin Pharmacol Ther. 84:362–369.

Roenneberg T, Wirz-Justice A, Merrow M. 2003. Life between clocks: daily temporal patterns of human chronotypes. J Biol Rhythms. 18:80–90.

Rutters F, Lemmens SG, Adam TC, Bremmer MA, Elders PJ, Nijpels G, Dekker JM. 2014. Is social jetlag associated with an adverse endocrine, behavioral, and cardiovascular risk profile? J Biol Rhythms. 29:377–383.

Silva GE, Goodwin JL, Sherrill DL, Arnold JL, Bootzin RR, Smith T, Walsleben JA, Baldwin CM, Quan SF. 2007. Relationship between reported and measured sleep times: the sleep heart health study (SHHS). J Clin Sleep Med. 3:622–630.

Somers VK, White DP, Amin R, Abraham WT, Costa F, Culebras A, Daniels S, Floras JS, Hunt CE, Olson LJ, et al. 2008. Sleep apnea and cardiovascular disease: an American Heart Association/American College of Cardiology Foundation Scientific Statement from the American Heart Association Council for High Blood Pressure Research Professional Education Committee, Council on Clinical Cardiology, Stroke Council, and Council on Cardiovascular Nursing. J Am Coll Cardiol. 52:686–717.

de Souza CM, Hidalgo MPL. 2014. Midpoint of sleep on school days is associated with depression among adolescents. Chronobiol Int. 31:199–205.

de Souza CM, Hidalgo MPL. 2015. The midpoint of sleep on working days: a measure for chronodisruption and its association to individuals’ well-being. Chronobiol Int. 32:341–348.

Sun X, Zheng B, Lv J, Guo Y, Bian Z, Yang L, Chen Y, Fu Z, Guo H, Liang P, et al. 2018. Sleep behavior and depression: Findings from the China Kadoorie Biobank of 0.5 million Chinese adults. J Affect Disord. 229:120–124.

Urbanek JK, Spira AP, Di J, Leroux A, Crainiceanu C, Zipunnikov V. 2017. Epidemiology of objectively measured bedtime and chronotype in US adolescents and adults: NHANES 2003-2006. Chronobiol Int.:1–19.

Vera B, Dashti HS, Gómez-Abellán P, Hernández-Martínez AM, Esteban A, Scheer FAJL, Saxena R, Garaulet M. 2018. Modifiable lifestyle behaviors, but not a genetic risk score, associate with metabolic syndrome in evening chronotypes. Sci Rep. 8:945.

Wittmann M, Dinich J, Merrow M, Roenneberg T. 2006. Social jetlag: misalignment of biological and social time. Chronobiol Int. 23:497–509.

Zee PC, Badr MS, Kushida C, Mullington JM, Pack AI, Parthasarathy S, Redline S, Szymusiak RS, Walsh JK, Watson NF. 2014. Strategic opportunities in sleep and circadian research: report of the Joint Task Force of the Sleep Research Society and American Academy of Sleep Medicine. Sleep. 37:219–227.

Zhai L, Zhang H, Zhang D. 2015. Sleep duration and depression among adults: a meta-analysis of prospective studies. Depress Anxiety. 32:664–670.

